# Quantification of compensatory motor neuron sprouting at the *Drosophila* larval neuromuscular junction

**DOI:** 10.64898/2026.09.06.749747

**Authors:** Lizzy Olsen, Aref Zarin

## Abstract

Following nervous system injury, surviving bystander neurons may rewire and form synapses with new postsynaptic targets to restore circuit function. This protocol quantifies compensatory motor neuron sprouting at the *Drosophila* larval neuromuscular junction. We describe larval dissection, immunostaining, Airyscan imaging, image masking, muscle-specific region-of-interest selection, and Fiji-based quantification of Dlg-positive area, bouton number, mean Dlg-positive area per bouton, NMJ arbor length, axonal versus NMJ branch points, ectopic muscle acquisition, and sprouting penetrance.

For complete details on the use and execution of this protocol, please refer to Olsen et al.^1^

**Graphical abstract:** 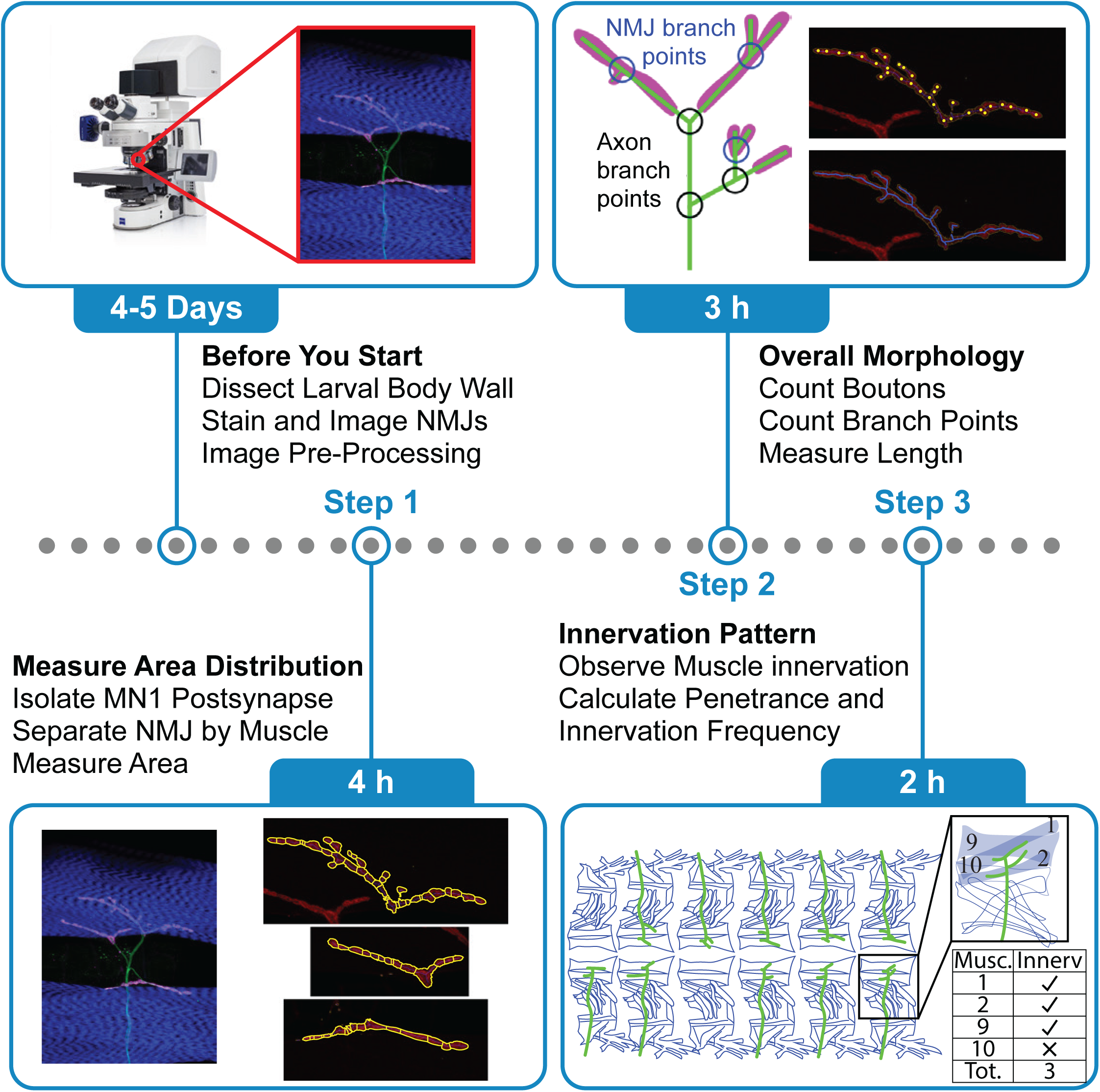

**Highlights:**

- Steps for larval dissection, immunostaining, Airyscan imaging, and preprocessing
- Instructions for Fiji-based masking to isolate single-neuron postsynaptic sites
- Guidance on muscle-specific quantification of synaptic area, boutons, and arbor length
- Steps for branch analysis to distinguish axonal sprouting from synaptic remodeling

**eTOC blurb:** Olsen and Zarin present a workflow for quantifying compensatory sprouting by a single motor neuron at the *Drosophila* larval neuromuscular junction. The protocol integrates Airyscan imaging, Fiji-based image masking, muscle-specific measurements, and branch classification to distinguish new axonal growth from remodeling of established synapses.

## Before you begin

This protocol quantifies the morphology and target distribution of a single genetically labeled motor neuron after neighboring motor neurons are ablated. In the associated study^1^, Olsen et al. used CQ-LexA and 27E09-LexA together to drive the apoptosis-inducing genes *reaper* and *head involution defective* in the tonic and phasic motor neurons that innervate dorsal longitudinal muscles 2, 9, and 10, thereby completely denervating these muscles. The surviving tonic motor neuron MN1 was selectively labeled using 94G06-Gal4-driven membrane-targeted GFP. Following denervation, MN1 forms ectopic neuromuscular junctions (NMJs) on the denervated muscles.

The protocol below describes how to quantify compensatory sprouting by MN1 onto the denervated dorsal longitudinal muscles. The workflow can also be adapted to other genetically labeled neurons and perturbations, provided that the axon of a single neuron can be distinguished unambiguously and an appropriate postsynaptic marker is available. The protocol measures Discs large (Dlg)-positive postsynaptic area as a proxy for NMJ area. It does not directly measure synaptic strength, active-zone number, or three-dimensional bouton volume.

## Innovation

Standard *Drosophila* NMJ morphometry focuses primarily on established synapses and commonly quantifies NMJ area, bouton number, bouton size, and arbor length^2,3^. The dual-driver motor-neuron ablation paradigm used by Olsen et al.^1^ completely removes tonic and phasic innervation from defined muscles while preserving and selectively labeling a neighboring tonic motor neuron, MN1. This design enables compensatory sprouting and ectopic innervation to be examined at single-neuron resolution.

Building on this paradigm, the protocol integrates several measurements to determine how MN1 changes its morphology and establishes ectopic innervation on denervated muscles. The labeled axon is used to mask the Dlg channel so that only postsynaptic structures associated with MN1 are retained. Muscle-specific regions of interest resolve the distribution of Dlg-positive area and boutons between the native and ectopically innervated targets. Branch points are classified as axonal or NMJ-associated based on their colocalization with Dlg and are further categorized by branch order. Whole-animal analysis quantifies muscle-specific innervation frequency, the number of muscles innervated, sprouting penetrance, and no-innervation penetrance. Unlike conventional NMJ morphometry, this integrated workflow distinguishes compensatory cross-neuronal sprouting and ectopic innervation of denervated muscles from remodeling or expansion of an established NMJ arbor.

## Install and configure the software

### Timing: less than 1 h

1. Install ZEN Lite from the [ZEISS download page](https://www.zeiss.com/microscopy/us/products/software/zeiss-zen-lite.html)
  a. Create or sign in to a ZEISS account if prompted.
  b. Install ZEN Lite with the default components and drivers.

**Note:** Airyscan processing requires an appropriate ZEN Blue license and is not available in ZEN Lite alone.

2. Install Fiji
  a. Download Fiji for your operating system from the [Fiji download page](https://imagej.net/software/fiji/downloads).
  b. Open the downloaded installer.

Follow the on-screen instructions to install Fiji.

3. Install the Drosophila NMJ Morphometrics macro in Fiji
  a. Download `Drosophila_NMJ_Morphometrics_EditsZarinLab.ijm` from the repository available in [Mendeley Data](https://data.mendeley.com/datasets/c9fdgs69xx/3).
  b. Copy the downloaded `.ijm` file into the Fiji `plugins` folder:
    i. On macOS, locate `Fiji.app`, right-click it, select **Show Package Contents,** and open the `plugins` folder.
    ii. On Windows, open the Fiji installation folder, typically `Fiji.app`, and then open the `plugins` folder.
  c. Restart Fiji and confirm that the Drosophila NMJ Morphometrics appears in the Plugins menu.
4. Increase the memory available to Fiji
  a. Select Edit > Options > Memory & Threads.
  b. Set **Maximum memory** to approximately 75% of the computer’s installed RAM.
  c. Restart Fiji.

**CRITICAL:** Default memory limitation is insufficient for opening the large images for quantification, so the maximum memory must be increased to 75%. However, do not allocate all available RAM to Fiji; reserve sufficient memory for the operating system and other required software, including ZEN. A computer with 16 GB of RAM or more is necessary to prevent Fiji from crashing.

### Prepare, dissect, and immunostain larvae

**Timing: 2–3 days** (excluding the time required for setting up crosses and rearing progeny)

5. Set up the required genetic crosses and rear the progeny under conditions appropriate for the experimental manipulation
  a. For constitutive *rpr*-mediated ablation, rear the crosses at 29°C to support efficient ablation^1^.
  b. Select late third-instar larvae by developmental stage and body size rather than chronological age because motor-neuron ablation can delay development.

**Note:** Include genotype-matched controls and process control and experimental samples in parallel. If feasible, randomize and blind samples before imaging and quantification.

6. Prepare a late third-instar larval fillet in HL3.1
  a. Place the larva ventral side up on a silicone elastomer dissection pad submerged in HL3.1.
  b. Pin the head and tail.
  c. Cut along the ventral midline with spring scissors.
  d. Remove the internal organs and tracheae without damaging the body-wall muscles or peripheral nerves.
  e. Place four additional pins to spread the body wall flat.

**CRITICAL:** Stretch the body wall sufficiently to pull the dorsal muscles taut, but do not tear the cuticle or distort the NMJs.

**Note:** To visualize the dorsal muscles, pin the larva ventral side up.

7. Fix and immunostain the fillet
  a. Remove HL3.1 and add enough 4% PFA to cover the entire fillet tissue. Fix for 10 min at 20°C–25°C.
  b. Remove the pins and transfer the fillet to a 0.5-mL microcentrifuge tube containing PBST. The used pins can be reused for future preparations.
  c. Wash three times in PBST for 10 min per wash on a rocker.
  d. Remove PBST and incubate the samples in blocking solution for 2 h at 20°C–25°C or 12–16 h at 4°C.
  e. Remove blocking solution and incubate the samples in primary-antibody solution for 2 h at 20°C–25°C or 12–16 h at 4°C.
  f. Wash three times in PBST for 10 min per wash.
  g. Incubate in secondary-antibody solution for 2 h at 20°C–25°C or 12–16 h at 4°C, protected from light.
  h. Wash three times in PBST for 10 min per wash, protected from light.

**CRITICAL:** Stain for Dlg to identify postsynaptic structures and label only one neuron strongly enough to trace its axon and terminals. Multiple labeled axons can invalidate masking and branch classification.

8. Mount three or four fillets per slide in Fluoromount-G with the muscle side facing the coverslip
  a. Align the cut midline parallel to the long axis of the slide.
  b. Store mounted slides at -20°C, protected from light, until imaging.

**Note:** Lining up the fillets with the midline parallel to the long side of the slide will limit the need for rotation during imaging and having the fillets close together will make it easier to find the next sample without changing objectives during imaging.

**Pause point:** Mounted slides can be stored at -20°C for several months, although imaging soon after staining minimizes fluorophore loss.

### Acquire high-resolution Airyscan NMJ images

#### Timing: 1–3 h per hemisegment for acquisition

9. Configure the ZEISS LSM 900 for Airyscan acquisition
  a. Enable Experiment Designer, Positions/Tiles, Z-stack, and Autosave in ZEN Blue.
  b. Use Smart Setup to add the GFP, Alexa Fluor 555, and CF647 channels, select the best signal configuration, and select Airyscan detection.
  c. Select bidirectional scanning, 16-bit acquisition, 0.8× zoom factor, and the Airyscan super-resolution (SR) frame-size preset.
  d. Use a 40× oil-immersion objective and the optimal z-interval, approximately 0.15 μm under the settings used by Olsen et al.^1^
  e. Record the pixel size, laser power, detector gain, z-interval, frame size, and bit depth, and save or export the complete Experiment Designer configuration.

**Note:** Use identical pixel size, z-interval, and bit depth for all samples that will be compared. Avoid saturated pixels.

**Note:** Using experiment designer and tiles functions, images can be taken from multiple animals on one slide overnight.

**Note:** Do not use Reuse to load settings from a previous .czi file acquired with Experiment Designer; this can cause ZEN to become unresponsive. Export and import the saved experiment configuration instead.

10. Identify suitable hemisegments
  a. Locate each larva with the 5× objective. Then switch to 20× objective.
  b. Screen the body wall with the 20× objective and exclude torn, folded, or severely distorted hemisegments.

**Note:** Center the stage clips before switching from 5x objective to 20x objective so that the objective cannot strike the slide holding clip.

11. Acquire the z-stack
  a. Center the selected hemisegment with the oil-immersion 40× objective and orient the ventral midline parallel to the upper edge of the image. Rotate opposite-side hemisegments by 180° when necessary
  b. Set channel-specific laser power and detector gain to capture the full signal range without saturation.
  c. Define the first and last z-positions so that the entire labeled axon and every associated Dlg-positive structure are included, with a small z-margin above and below the visible signal. Choose the optimal z-interval, approximately 0.15 μm under the settings used by Olsen et al.^1^
  d. Move to the center of the z-stack before adding the position to Experiment Designer.
  e. Stop live scanning before duplicating the configured experiment; duplicating during live scanning can reset the frame size to 512 × 512 pixels.
  f. Repeat steps 11a–11e for two hemisegments on each animal on the slide. Verify the frame size, z-center, channels, and autosave location for every position.
  g. Start acquisition.

**CRITICAL:** Keep laser power and detector gain as consistent as possible across samples in the same experiment. A shifted or truncated z-stack biases area and length measurements (Troubleshooting, problem 1). Ensure that the SR preset is selected and that the frame size remains set to SR.

**Note:** Center the stage clips before switching from 20x objective to 40x objective so that the objective cannot strike the slide holding clip.

**Note:** Image two analyzable hemisegments from each larva when possible and acquire at least 12 hemisegments from at least six larvae per condition. Increase the sample size when the expected effect is small or variable.

### Image pre-processing

#### Timing: Approximately 20 min per hemisegment for preprocessing

12. Process the raw z-stacks in ZEN Blue
  a. Open Processing, switch from Single to Batch, and select Airyscan Processing.
  b. Add the raw Airyscan .czi z-stacks for samples acquired with the same settings.

**Optional:** Select all files, clear Use Input Folder as Output Folder, and assign an output folder so that the post processing files are not saved in individual subfolders.

Under Method Parameters, select 3D Processing and Auto Filter for one file.
Copy and paste the parameters between files; changing the parameters for one file does not automatically update the others.
Select Apply and allow every file to finish processing.

**Note:** At the end of this step, the output folder should contain one Airyscan-processed z-stack for every hemisegment. Retain the original raw .czi files.

13. Export the Dlg channel as individual 8-bit TIFF slices
  a. In Processing, switch to Batch, select Image Export, and add the Airyscan-processed z-stacks.
  b. Select all files, clear Use Input Folder as Output Folder, and assign a dedicated parent quantification folder.
  c. Select Show All to display the complete export settings.
  d. Set File Type to TIFF and select Convert to 8-bit.
  e. Select Apply Display Curve and Channel Color. To change the display curve or channel color, open each file outside of batch processing, adjust the minimum and maximum pixel values or colors, and save the file before exporting. Ensure that the display curve is consistent across all files.
  f. Select Individual Channel Images and Short Format. Clear Merged Channel Image and Burn-in Annotation.
  g. Under Define Subset > Channels, retain only the channel containing Dlg, such as Alexa Fluor 555.
  h. Select Create Folder so that the TIFF slices from each hemisegment are exported into a separate subfolder.
  i. Copy the complete export settings to every file, select Apply, and allow all slices to export.

**Note:** The parent quantification folder should contain one uniquely named subfolder per hemisegment, with the individual Dlg-channel TIFF slices from that z-stack.

**CRITICAL:** Do not burn scale bars, labels, or other annotations into images used for quantification; these elements alter image pixels and can interfere with thresholding and area measurements.

14. Convert the Dlg slices into a stack and maximum-intensity projection in Fiji
  a. Open Fiji and start the Drosophila NMJ Morphometrics plugin.
  b. Set Stack identifier to stack and Projection identifier to flatstack. (Figure 1A)
  c. Select Convert to Stack and click OK.
  d. Select the parent quantification folder containing the hemisegment subfolders.
  e. Allow the plugin to assemble the slices into a stack, generate a maximum-intensity projection, and save both files in the corresponding subfolder.
  f. Confirm that each subfolder contains one TIFF stack with a filename beginning with stack and one maximum-intensity projection beginning with flatstack.

**Figure 1.**
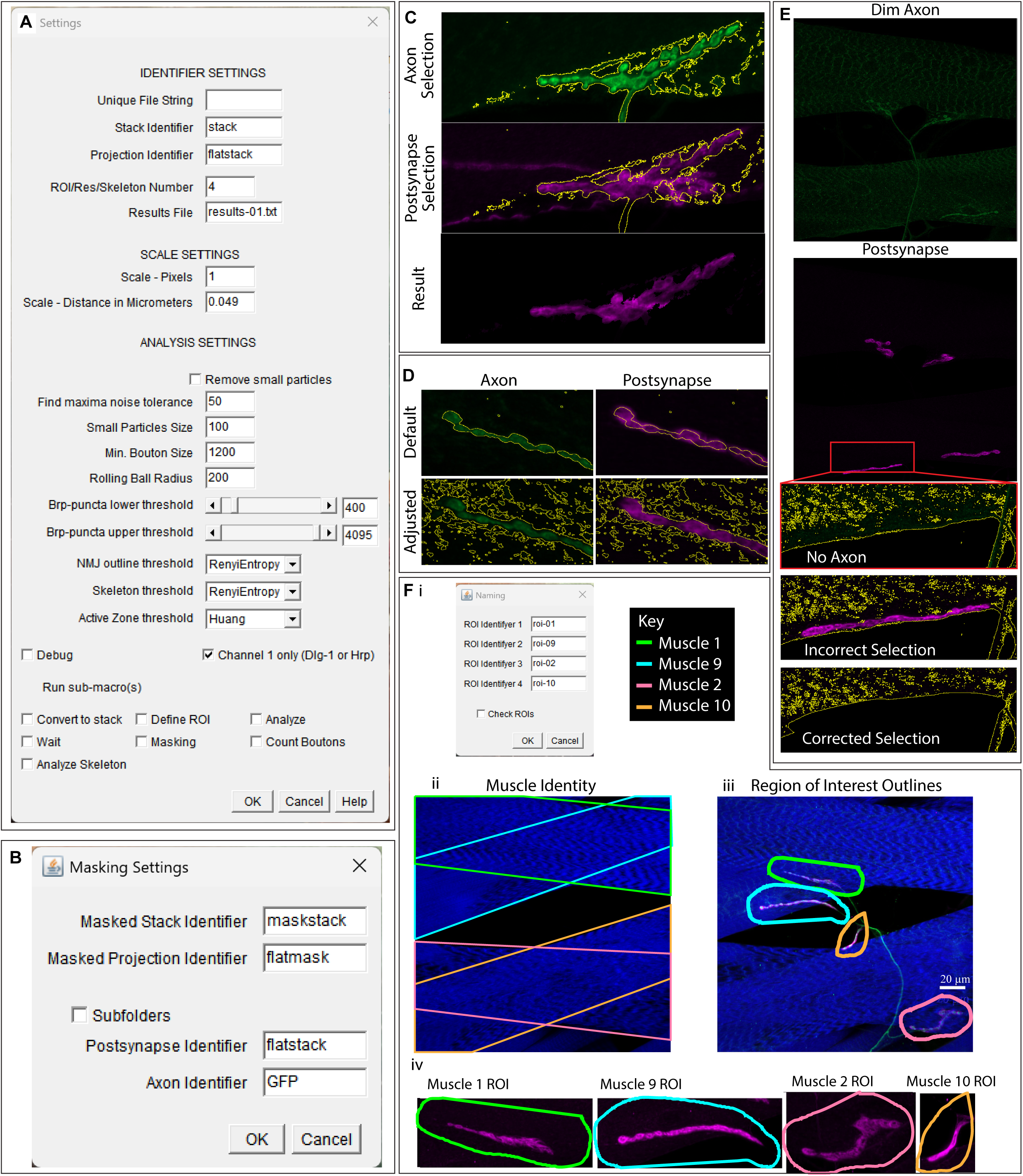
Isolating postsynaptic structures associated with the labeled axon. Screenshots of the macro main dialogue box (A) and masking dialogue box (B). (C) A threshold-based selection of the labeled axon is transferred to the Dlg channel to remove non-colocalized Dlg signal. (D) The default axon threshold can exclude associated Dlg signal; expanding the selection retains the complete postsynaptic structure. (E) When axonal fluorescence is dim, a permissive threshold can retain non-colocalized Dlg, which is removed manually. (F) (i) Screenshot of the Naming dialogue box opened after selecting Define ROI. Muscle identities (ii) guide muscle-specific ROI placement (iii–iv). Muscle 1 is green, muscle 2 is pink, muscle 9 is cyan, and muscle 10 is orange.

**Note:** Opening the slices, assembling the stack, and generating the projection can take several minutes per hemisegment.

**Pause point:** After confirming that the stack and flatstack files were generated correctly, compress the individual exported Dlg slices into a ZIP archive. Verify the archive before deleting uncompressed working copies. Retain the raw .czi files, Airyscan-processed .czi files, TIFF stack, and maximum-intensity projection.

15. Export the labeled-axon channel as a maximum-intensity projection

a. Open an Airyscan-processed z-stack in ZEN.
b. In Processing, select Orthogonal Projection.
c. Set Projection Plane to XY and Method to Maximum.
d. Set the start position to the first slice and the thickness or end position to include the complete z-stack.
e. Select Apply and save the resulting projection .czi file.
f. Repeat steps 15a–15e for every hemisegment.
g. Use Image Export to export only the labeled-axon channel, such as Alexa Fluor 488, from each projection as an 8-bit TIFF.
h. Use the applicable export settings in step 13, but clear Create Folder because each projection contains only one image.
i. Prefix each exported axon projection with GFP_ and remove the automatically generated orthogonal-projection suffix so that the portion of the filename following GFP_ exactly matches its corresponding hemisegment subfolder.
j. Move each GFP_ projection into its corresponding hemisegment subfolder.

**Note:** Orthogonal Projection is not available in Batch mode and must be applied to each hemisegment individually.

**CRITICAL:** Before beginning quantification, confirm that every hemisegment subfolder contains exactly one Dlg stack, one Dlg maximum-intensity projection, and one matching labeled-axon projection (Troubleshooting, problem 5).

Key resources table.

Key resources table
| REAGENT or RESOURCE | SOURCE | IDENTIFIER |
| --- | --- | --- |
| **Antibodies** |  |  |
| Rabbit polyclonal anti-GFP | Invitrogen | Cat. #A-11122; RRID: AB_221569 |
| Mouse monoclonal anti-Dlg | Developmental Studies Hybridoma Bank | RRID: AB_528203 |
| Alexa Fluor 488 goat anti-rabbit IgG | Invitrogen | Cat. #A-11034; RRID: AB_2576217 |
| Alexa Fluor Plus 555 goat anti-mouse IgG | Invitrogen | Cat. #A32727; RRID: AB_2633276 |
| **Chemicals, peptides, and recombinant proteins** |  |  |
| CF647 phalloidin | Biotium | Cat. #00041 |
| Paraformaldehyde | Any reputable supplier | n/a |
| Normal goat serum | Any reputable supplier | n/a |
| Normal donkey serum | Any reputable supplier | n/a |
| Phosphate-buffered saline | Any reputable supplier | n/a |
| Triton X-100 | Any reputable supplier | n/a |
| Fluoromount-G | SouthernBiotech | Cat. #0100-01 |
| **Experimental models: Organisms/strains** |  |  |
| D. melanogaster: w; 94G06-Gal4 | Bloomington Drosophila Stock Center <sup>4,5</sup> | BDSC #40701 |
| D. melanogaster: w; CQ-LexA | Cooney et al.<sup>6</sup> Sitaula et al.<sup>7</sup> | n/a |
| D. melanogaster: w; 27E09-LexA | Olsen et al.<sup>1</sup> | n/a |
| D. melanogaster: w; CQ-LexA, 27E09-LexA | Olsen et al.<sup>1</sup> | n/a |
| D. melanogaster: w; CQ-LexA, 27E09-LexA; 94G06-Gal4 | Olsen et al.<sup>1</sup> | n/a |
| D. melanogaster: w[*]; P{y[+t7.7] w[+mC]=13xlexAop2-rpr.H}su(Hw)attP5/CyO; Dr[1]/TM6C, Sb[1] | Robin Harris; Bloomington Drosophila Stock Center | BDSC #605696, as reported in Olsen et al.<sup>1</sup> |
| D. melanogaster: 10×UAS-myr::GFP | Bloomington Drosophila Stock Center | BDSC #32198 |
| **Software and algorithms** |  |  |
| Fiji | ImageJ | https://imagej.net/software/fiji/downloads |
| ZEN Blue | ZEISS | https://www.zeiss.com/microscopy/us/products/software/zeiss-zen.html |
| ZEN Blue | ZEISS | https://www.zeiss.com/microscopy/us/products/software/zeiss-zen-lite.html |
| Drosophila NMJ Morphometrics | Schenck laboratory<sup>2,3</sup> | https://figshare.com/articles/dataset/Drosophila_NMJ_Morphometrics/2077399/1 |
| Drosophila_NMJ_Morphometrics_EditsZarinLab.ijm |  | https://data.mendeley.com/datasets/c9fdgs69xx/3<br>DOI:10.17632/c9fdgs69xx.3 |
| R and RStudio, optional | R Foundation; Posit | https://posit.co/download/rstudio-desktop/ |
| **Other** |  |  |
| LSM 900 confocal microscope with Airyscan and 40× oil objective | ZEISS | LSM 900 |
| Stereo microscope | ZEISS or equivalent | Stemi 305 or equivalent |
| Silicone elastomer dissection pad | Dow or equivalent | Sylgard 184 or equivalent |
| Spring scissors and fine insect pins | Fine Science Tools or equivalent | n/a |

## Materials and equipment

### HL3.1, 1 L

Materials and equipment HL3.1, 1 L
| Reagent | Final concentration | Amount for 1 L |
| --- | --- | --- |
| NaCl | 70 mM | 4.09 g |
| KCl | 5 mM | 0.373 g |
| CaCl<sub>2</sub> | 1.5 mM | 0.166 g anhydrous CaCl<sub>2</sub> or 0.221 g CaCl<sub>2</sub>·2H<sub>2</sub>O |
| MgCl<sub>2</sub> | 4 mM | 0.381 g anhydrous MgCl<sub>2</sub> or molar equivalent of hydrate |
| NaHCO<sub>3</sub> | 10 mM | 0.840 g |
| Trehalose | 5 mM | 1.89 g trehalose dihydrate |
| Sucrose | 115 mM | 39.35 g |
| HEPES | 5 mM | 1.19 g |
| ddH<sub>2</sub>O | n/a | Bring to 1 L |

Adjust to pH 7.2. Store at 4°C for up to 1 month. Warm an aliquot to 20°C–25°C before dissection. This formulation follows Feng et al.^8^

### PBST, 1 L

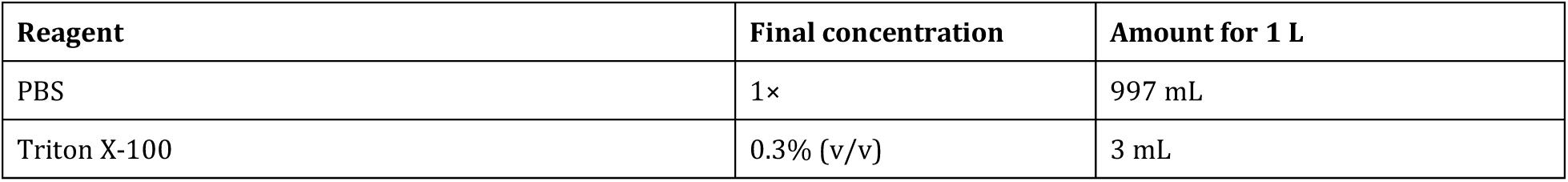

Store at 20°C–25°C for up to 6 months.

### 4% PFA, 40 mL

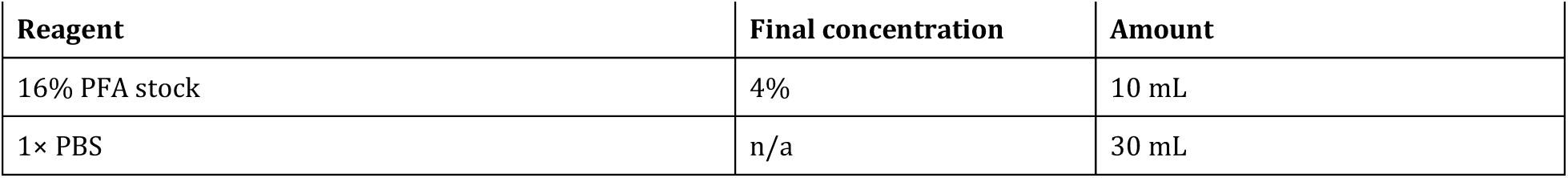

Aliquot and store at -20°C for up to 6 months. Thaw an aliquot immediately before use and do not repeatedly freeze and thaw it.

**CRITICAL:** PFA is toxic. Prepare and handle PFA in a certified chemical fume hood with appropriate personal protective equipment and dispose of it according to institutional requirements.

### Blocking solution, 40 mL

| Reagent | Final concentration | Amount |
| --- | --- | --- |
| Normal goat serum | 2.5% (v/v) | 1 mL |
| Normal donkey serum | 2.5% (v/v) | 1 mL |
| PBST | n/a | 38 mL |

Store at 4°C for up to 1 month.

### Primary-antibody solution, 300 μL

| Reagent | Final concentration | Amount |
| --- | --- | --- |
| Blocking solution | n/a | 296.4 µL |
| Rabbit anti-GFP | 1:500 | 0.6 µL |
| Mouse anti-Dlg | 1:100 | 3 µL |

Prepare immediately before use. Scale the volume to cover all samples in one tube.

### Secondary-antibody solution, 300 μL

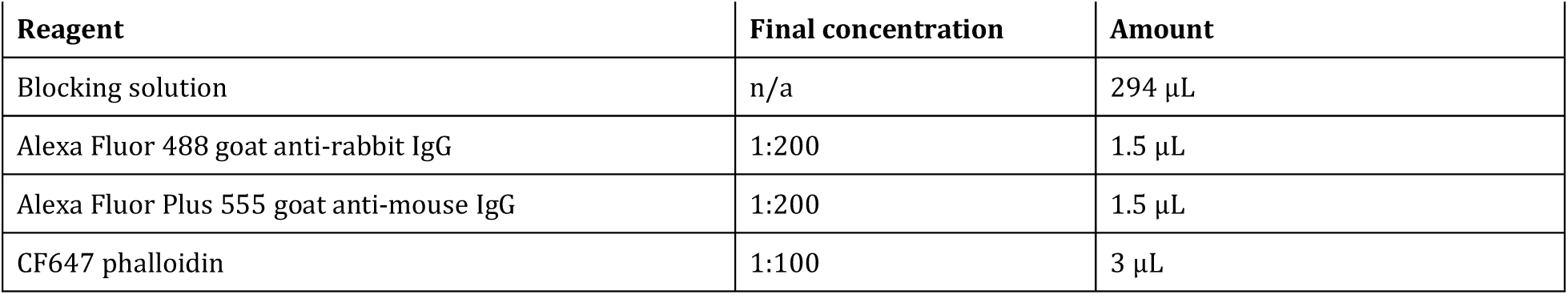

Secondary-antibody solution, 300 μL
| Reagent | Final concentration | Amount |
| --- | --- | --- |
| Blocking solution | n/a | 294 μL |
| Alexa Fluor 488 goat anti-rabbit IgG | 1:200 | 1.5 μL |
| Alexa Fluor Plus 555 goat anti-mouse IgG | 1:200 | 1.5 μL |
| CF647 phalloidin | 1:100 | 3 μL |

Prepare immediately before use and protect from light.

**Alternatives:** Change the primary and secondary antibodies when the neuron of interest is labeled with a fluorophore other than GFP. Preserve spectral separation among the labeled axon, Dlg, and phalloidin channels.

### Recommended image-acquisition and preprocessing settings

Settings are grouped by acquisition and preprocessing stage for the ZEISS LSM 900 Airyscan workflow described in the protocol. Instrument-dependent values are starting points and must be optimized without saturation.

| Dialogue box or workflow stage | Parameter | Setting | Purpose or caution |
| --- | --- | --- | --- |
| **Image acquisition** | Objective | 40× oil immersion | Used for high-resolution NMJ acquisition after screening the specimen at lower magnification. |
| Image acquisition | Experiment Designer | Enabled | Use for multiposition or overnight acquisition; also verify that Autosave is enabled. |
| Image acquisition | Positions/Tiles | Enabled; use Positions for individual hemisegments | Tile regions are an alternative for whole-animal fields that do not fit within one frame. |
| Image acquisition | Z-stack | Enabled | Required to capture the entire labeled axon and all associated Dlg-positive structures. |
| Image acquisition | Channels | GFP, Alexa Fluor 555, and CF647 | Use Smart Setup, select the best signal configuration, and choose Airyscan detection. |
| Image acquisition | Digital zoom | 0.8× | Captures the dorsal muscle field under the reported setup. |
| Image acquisition | Scan-area orientation | Rotate opposite-side hemisegments by 180° when needed | Place the ventral midline parallel to the upper edge of the image. |
| Image acquisition | Frame size | Airyscan super-resolution preset | Stop live scanning before duplicating an experiment and verify that the frame size has not reset to 512 × 512 pixels. |
| Image acquisition | Scan direction | Bidirectional | Reduces acquisition time. |
| Image acquisition | Bit depth | 16 bit | Preserves the intensity range; use the same bit depth for all compared samples. |
| Image acquisition | Pixel size | Record from the acquisition metadata | Keep identical within an experiment and do not assume the value for a different configuration. |
| **Channels and signal settings** | Laser power: CF647/phalloidin | Approximately 0.01% | Reported starting value; optimize for the specimen and instrument without saturating pixels. |
| Channels and signal settings | Laser power: Dlg | Approximately 0.02%–0.04% | Reported starting range; record the final value and apply it consistently within the experiment. |
| Channels and signal settings | Laser power: labeled axon/GFP | Approximately 0.25%–1.5% | Reported starting range; optimize to trace the complete axon while avoiding saturation. |
| Channels and signal settings | Detector master gain | Approximately 750; approximately 650 for bright phalloidin signal | Instrument- and sample-dependent; record the final channel-specific values. |
| **Experiment Designer and positions** | Add position (+) | Add only after moving to the center of the z-stack | Prevents a shifted or truncated z-stack during multiposition acquisition. |
| Experiment Designer and positions | Duplicate | Duplicate the configured experiment for additional positions | Set laser power and gain first, stop live scanning, and verify the frame size, channels, z-center, and autosave location after duplication. |
| Experiment Designer and positions | Experiment configuration | Export and import the saved configuration | Do not use Reuse to load settings from an earlier .czi acquired with Experiment Designer because ZEN can become unresponsive. |
| Experiment Designer and positions | Autosave folder | User-defined; verify for every position | Choose a location with sufficient space and do not overwrite prior acquisitions. |
| Experiment Designer and positions | Image name | Include genotype and slide number | Use short, unique names that remain traceable through export and quantification. |
| **Z-stack** | First position | Shallowest point at which the labeled axon or associated Dlg signal is visible | Include a small z-margin beyond the visible signal. |
| Z-stack | Last position | Deepest point at which the labeled axon or associated Dlg signal is visible | Include a small z-margin beyond the visible signal. |
| Z-stack | Interval | Optimal; approximately 0.15 $\mu\text{m}$ with the reported 40× settings | Use the same z-interval for all samples being compared. |
| Z-stack | Number of slices | Automatically determined by the first/last range and interval | Do not set independently unless the acquisition configuration requires it. |
| Z-stack | Center position | Move to the midpoint before adding the Experiment Designer position | Confirm that the z-stack remains centered for every hemisegment. |
| **Airyscan processing** | Processing mode | Batch | Process together only z-stacks acquired with the same settings. |
| Airyscan processing | 3D Processing | Selected | Uses information from adjacent z-slices during Airyscan processing. |
| Airyscan processing | Auto Filter | Selected | Apply identically to every file in the batch. |
| Airyscan processing | Output folder | Dedicated folder; Use Input Folder as Output Folder cleared | Retains the raw .czi files and prevents them from being overwritten. |
| **Dlg-channel image export** | Processing mode | Batch; Show All selected | Displays the complete export settings and applies them consistently across the batch. |
| Dlg-channel image export | Output folder | Dedicated parent quantification folder; Use Input Folder as Output Folder cleared | Keeps exported images separate from Airyscan-processed .czi files. |
| Dlg-channel image export | File type | TIFF; Convert to 8-bit selected | Exports images in the format required for Fiji processing. |
| Dlg-channel image export | Apply Display Curve and Channel Color | Selected | Set a nonsaturating range that retains dim Dlg signal, record its limits, and apply it unchanged to all samples in the experiment. |
| Dlg-channel image export | Individual Channel Images | Selected | Exports the Dlg channel as separate images rather than as a merged multichannel image. |
| Dlg-channel image export | Short Format | Selected | Produces concise filenames compatible with downstream organization. |
| Dlg-channel image export | Merged Channel Image | Cleared | A merged image is not used for quantification. |
| Dlg-channel image export | Burn-in Annotation | Cleared | Do not burn scale bars, labels, or other annotations into images used for quantification. |
| Dlg-channel image export | Define Subset > Channels | Dlg channel only; for example, Alexa Fluor 555 | Excludes the other fluorescence channels from this export. |
| Dlg-channel image export | Create Folder | Selected | Creates one uniquely named subfolder per hemisegment containing the individual Dlg TIFF slices. |
| **Labeled-axon projection and export** | Projection Plane | XY | Generates a frontal projection corresponding to the analysis view. |
| Labeled-axon projection and export | Projection Method | Maximum | Generates a maximum-intensity projection of the labeled axon. |
| Labeled-axon projection and export | Start position | First slice | Include the complete z-stack. |
| Labeled-axon projection and export | Thickness/end position | Complete number or range of slices in the z-stack | Do not truncate the labeled axon or terminals. |
| Labeled-axon projection and export | Image-export file type | TIFF; Convert to 8-bit selected | Exports the axon projection for use in Fiji masking and branch classification. |
| Labeled-axon projection and export | Apply Display Curve and Channel Color | Selected | Use a nonsaturating range that preserves dim axon branches and record the applied limits. |
| Labeled-axon projection and export | Individual Channel Images and Short Format | Selected | Export only the required axon image with a concise filename. |
| Labeled-axon projection and export | Merged Channel Image and Burn-in Annotation | Cleared | Prevents merged or annotated pixels from entering the analysis. |
| Labeled-axon projection and export | Define Subset > Channels | Labeled-axon channel only; for example, Alexa Fluor 488 | Exports only the axon channel from the projection. |
| Labeled-axon projection and export | Create Folder | Cleared | Each projection contains one image; move the exported GFP_ file into the matching hemisegment subfolder. |

*Note: Generic interface controls such as Delete, folder-navigation buttons, and stage-movement shortcuts are omitted because they do not define acquisition or processing parameters. Record the complete Experiment Designer configuration and the final channel-specific laser power, gain, display range, pixel size, and z-interval for each experiment.*

### NMJ Morphometrics settings used in this workflow

Settings are grouped by dialogue box and workflow stage. Values apply to the single-channel Dlg/axon-masking workflow described in the protocol.

| Dialogue box or workflow stage | Parameter | Setting | Purpose or caution |
| --- | --- | --- | --- |
| Main dialogue box: baseline settings | Unique file string | Leave blank | Optional filter; use only when files must be restricted by a unique filename string. |
| Main dialogue box: baseline settings | ROI/Result/Skeleton number | 4 | One entry for each analyzed muscle: 1, 2, 9, and 10. |
| Main dialogue box: baseline settings | Scale-Pixels | 1 | Enables calibrated measurements in micrometers rather than pixels. |
| Main dialogue box: baseline settings | Scale—distance in micrometers | Read from the .czi metadata; approximately 0.049 $\mu\text{m}$/pixel in the reported setup | Do not assume this value for a different acquisition configuration. |
| Main dialogue box: baseline settings | Remove small particles | False | Avoids deleting genuine small Dlg-positive structures. |
| Main dialogue box: baseline settings | Minimum bouton size | 1,200 pixels | Plugin setting; boutons are counted manually in this workflow. |
| Main dialogue box: baseline settings | Rolling-ball radius | 200 pixels | Starting value for background subtraction; adjust only after inspecting the output. |
| Main dialogue box: baseline settings | NMJ-outline threshold | Rényi entropy | Starting threshold; inspect every generated outline. |
| Main dialogue box: baseline settings | Skeleton threshold | Rényi entropy | Starting threshold; inspect and clean every generated skeleton. |
| Main dialogue box: baseline settings | Channel 1 only (Dlg-1 or Hrp) | True | Disables Brp/active-zone analysis, which is not performed in this workflow. |
| Main dialogue box: baseline settings | Debug | False | Enable only for troubleshooting. |
| **Convert exported Dlg slices to stacks and projections** | Stack identifier | stack | Prefix assigned to the assembled Dlg z-stack. |
| Convert exported Dlg slices to stacks and projections | Projection identifier | flatstack | Prefix assigned to the Dlg maximum-intensity projection. |
| **Masking dialogue box** | Stack identifier | stack | Identifies the input Dlg stack before masking. |
| Masking dialogue box | Projection identifier | flatstack | Identifies the input Dlg projection before masking. |
| Masking dialogue box | Masked stack identifier | maskstack | Prefix assigned to the masked Dlg stack. |
| Masking dialogue box | Masked projection identifier | flatmask | Prefix assigned to the masked Dlg projection. |
| Masking dialogue box | Subfolders | False | Use when the matched Dlg and GFP files are stored together within each hemisegment folder. If channels are stored in separate folders, enter the corresponding folder names instead. |
| Masking dialogue box | Postsynapse identifier | flatstack | Identifies the unmasked Dlg projection used as the masking input. |
| Masking dialogue box | Axon identifier | GFP | Identifies the matching labeled-axon projection; filenames begin with GFP_. |
| **Define ROIs dialogue box** | Stack identifier | maskstack | Uses the masked Dlg stack. |
| Define ROIs dialogue box | Projection identifier | flatmask | Uses the masked Dlg projection. |
| Define ROIs dialogue box | ROI identifiers | roi-01, roi-02, roi-09, and roi-10 | One ROI identifier for each muscle analyzed. |
| Define ROIs dialogue box | Check ROIs | False for the initial analysis; True when reviewing saved ROIs | Inspect and adjust existing ROIs when this option is selected. |
| Define ROIs dialogue box | No ROI | False by default; select for an uninnervated muscle | Make a temporary selection before choosing No ROI so that the plugin advances without saving an ROI. |
| **Analyze Dlg-positive area** | Stack identifier | maskstack | Uses the masked Dlg stack. |
| Analyze Dlg-positive area | Projection identifier | flatmask | Uses the masked Dlg projection. |
| Analyze Dlg-positive area | ROI/Result/Skeleton number | 4 | Processes muscles 1, 2, 9, and 10. |
| Analyze Dlg-positive area | ROI identifiers | roi-01, roi-02, roi-09, and roi-10 | Must match the saved muscle-specific ROI files. |
| Analyze Dlg-positive area | Result identifiers | res-01, res-02, res-09, and res-10 | Assigns a separate result output to each ROI. |
| Analyze Dlg-positive area | Rolling-ball radius | 200 pixels | Starting value; increase if background subtraction removes genuine boutons. |
| Analyze Dlg-positive area | NMJ-outline threshold | Rényi entropy | Reanalyze only affected ROIs with a documented alternative threshold when necessary. |
| Analyze Dlg-positive area | Skeleton threshold | Rényi entropy | Starting threshold used to generate the skeleton for later inspection and correction. |
| **Count Boutons/Branching dialogue box** | Stack identifier | maskstack | Uses the masked Dlg stack. |
| Count Boutons/Branching dialogue box | Projection identifier | flatmask | Uses the masked Dlg projection. |
| Count Boutons/Branching dialogue box | ROI identifiers | roi-01, roi-02, roi-09, and roi-10 | Counts boutons and NMJ branch points separately within each muscle ROI. |
| Count Boutons/Branching dialogue box | Subfolders | False | The Dlg and GFP files are stored together within each hemisegment folder in this workflow. |
| Count Boutons/Branching dialogue box | Postsynapse identifier | flatmask | Identifies the masked Dlg projection. |
| Count Boutons/Branching dialogue box | Axon identifier | GFP | Identifies the matching labeled-axon projection. |
| Count Boutons/Branching dialogue box | Count Branching | True | Enables classification of axonal and NMJ-associated branch points by branch order. |
| Count Boutons/Branching dialogue box | Results-file name | User-defined; for example, boutons_and_branchpoints.txt | Use a unique filename to avoid overwriting an existing analysis. |
| Count Boutons/Branching dialogue box | Read File | False for a new analysis; True when reviewing a saved analysis | When reviewing, load the matching results file and inspect the saved bouton points. |
| Analyze Skeleton dialogue box | Stack identifier | maskstack | Uses the masked Dlg stack. |
| Analyze Skeleton dialogue box | Projection identifier | flatmask | Uses the masked Dlg projection. |
| Analyze Skeleton dialogue box | Results-file name | User-defined; for example, NMJ_skeleton_results.txt | Use a unique filename to avoid overwriting an existing analysis. |
| Analyze Skeleton dialogue box | Result identifiers | res-01, res-02, res-09, and res-10 | Must match the ROI-specific result images generated during analysis. |
| Analyze Skeleton dialogue box | Skeleton identifiers | skeleton-01, skeleton-02, skeleton-09, and skeleton-10 | Assigns one corrected skeleton file to each muscle ROI. |
| Analyze Skeleton dialogue box | Check Skeleton | False for the initial analysis; True when reviewing corrected skeletons | Reopens previously corrected skeletons for inspection. |

*Note: Find Maxima noise tolerance, Brp-puncta thresholds, and the active-zone threshold are omitted because Channel 1 only is set to True and active zones are not quantified. Small-particle size is also omitted because Remove small particles is set to False. First-, second-, and third-order branch fields are outputs rather than preset parameters.*

## Step-by-step method details

**Note:** The ImageJ macro processes all exported TIFF stacks in a folder. For each condition, quantify at least twelve hemisegments from six animals.

### Isolate the postsynaptic structures associated with the labeled axon

#### Timing: approximately 5 min per image

This step uses the labeled-axon projection to mask the Dlg stack and projection, thereby retaining only Dlg-positive structures associated with the neuron of interest (Figure 1A-E).

1. Configure the masking routine. **(Figure 1A and 1B)**

a. Open the NMJ Morphometrics plugin.
b. Set Stack identifier to stack and Projection identifier to flatstack.
c. Select **Masking** and click **OK**.
d. Set Masked stack identifier to maskstack and Masked projection identifier to flatmask.
e. Set the postsynapse identifier to flatstack and the axon identifier to GFP, or enter the corresponding subfolder names if the files are organized in separate channel folders.
f. Select the parent quantification folder containing subfolders to analyze.

**Note:** The masking routine will not start unless it detects the same number of stack, flatstack, and GFP_ files. If no images open, verify the filenames and identifiers (Troubleshooting, problem 5).

2. Define the axon mask for each hemisegment

a. In the axon-projection window, adjust the lower brightness threshold until the selection covers the complete labeled axon and a narrow margin surrounding it.
b. Convert the thresholded signal to a selection.

**Note:** Do not click OK in the plugin dialog until a selection is present in the Dlg-projection window.

Switch to the window with the postsynapse tiff (Dlg-projection window) and restore the selection (Edit > Selection > Restore Selection).
Confirm that the selection includes all Dlg-positive structures opposed to the labeled axon and excludes Dlg associated with other axons (**Figure 1C and 1D**).
Manually add missing regions or remove non-colocalized Dlg when a dim axon prevents a single threshold from producing an accurate mask (**Figure 1E**; Troubleshooting, problem 2).
Click **OK** and allow the plugin to save the masked stack and projection.

**Note:** Do not close the image windows while the plugin is processing them. At the end of this step, each hemisegment subfolder should contain new files beginning with maskstack and flatmask.

**CRITICAL:** Do not rely on automatic thresholding without visual verification. An overly restrictive mask underestimates NMJ area, whereas an overly permissive mask includes postsynaptic structures from other neurons.

### Define muscle-specific regions of interest

#### Timing: approximately 5 min per image

This step assigns each retained postsynaptic structure to dorsal muscles 1, 2, 9, or 10 (**Figure 1F**).

3. Draw and save four muscle-specific ROIs
  a. Open the NMJ Morphometrics plugin.
  b. Set **Stack identifier** to maskstack, **Projection identifier** to flatmask, and **ROI/Result/Skeleton number to 4**.
  c. Select **Define ROI** option and click OK.
  d. Enter roi-01, roi-02, roi-09, and roi-10 as the ROI identifiers.
  e. Select the parent quantification folder.
  f. For each requested muscle, use the freehand tool to enclose all associated Dlg-positive structures without including a structure from another muscle.
  g. When the neuron does not innervate the requested muscle, make a temporary selection and select **No ROI** so that the plugin advances without saving an ROI.

**CRITICAL:** Maximum-intensity projections can make structures on different z-planes appear colocalized. To avoid this mistake and to accurately assign structures to muscles, open the full z-stack, the axon projection, and the phalloidin channel in ZEN Blue.

**Note:** The body-wall membrane can pull away from the phalloidin-labeled actin during dissection, so an associated Dlg-positive structure may not completely overlap the phalloidin signal. Use its z-position and the trajectory of the labeled axon when assigning the muscle.

**Note:** If ROIs already exist, select **Check ROIs**, inspect each saved ROI, and adjust it if needed.

**Note:** At the end of this step, each hemisegment subfolder should contain one saved ROI file for every innervated muscle. Do not create an ROI file for a muscle scored as No ROI.

### Quantify Dlg-positive NMJ area on each muscle

#### Timing: approximately 2–5 min per image after setup

This step performs background subtraction and thresholding within each muscle-specific ROI and reports calibrated Dlg-positive area (**Figure 2A-E**).

1. Configure and run the area analysis (**Figure 2A, panels i and ii**)
2. Open the NMJ Morphometrics plugin.
3. Set **Stack identifier** to maskstack and **Projection identifier** to flatmask.
4. Enter a unique results-file name, such as NMJ_area_results.txt.
5. Set **Scale pixels** to 1 and enter the pixel size recorded in the .czi metadata.
6. Set **ROI/Result/Skeleton number** to 4 and **Rolling-ball radius** to 200 pixels.
7. Set both the NMJ-outline and skeleton thresholds to **RenyiEntropy**.
8. Select **Analyze** and **Wait** options, enter the matching ROI and result identifiers, and choose the parent quantification folder.
9. Inspect the outline generated for every ROI (**Figure 2B and 2C**).
10. If the outline omits genuine Dlg-positive area, test a more inclusive method such as Huang or Triangle (**Figure 2D**). If the outline includes background or non-NMJ signal, test a more restrictive method such as Li (**Figure 2E**). Troubleshooting, problem 3).
11. ii. Reanalyze only the inaccurate ROI or image with the selected alternative threshold and record the method used.

**Figure 2.**
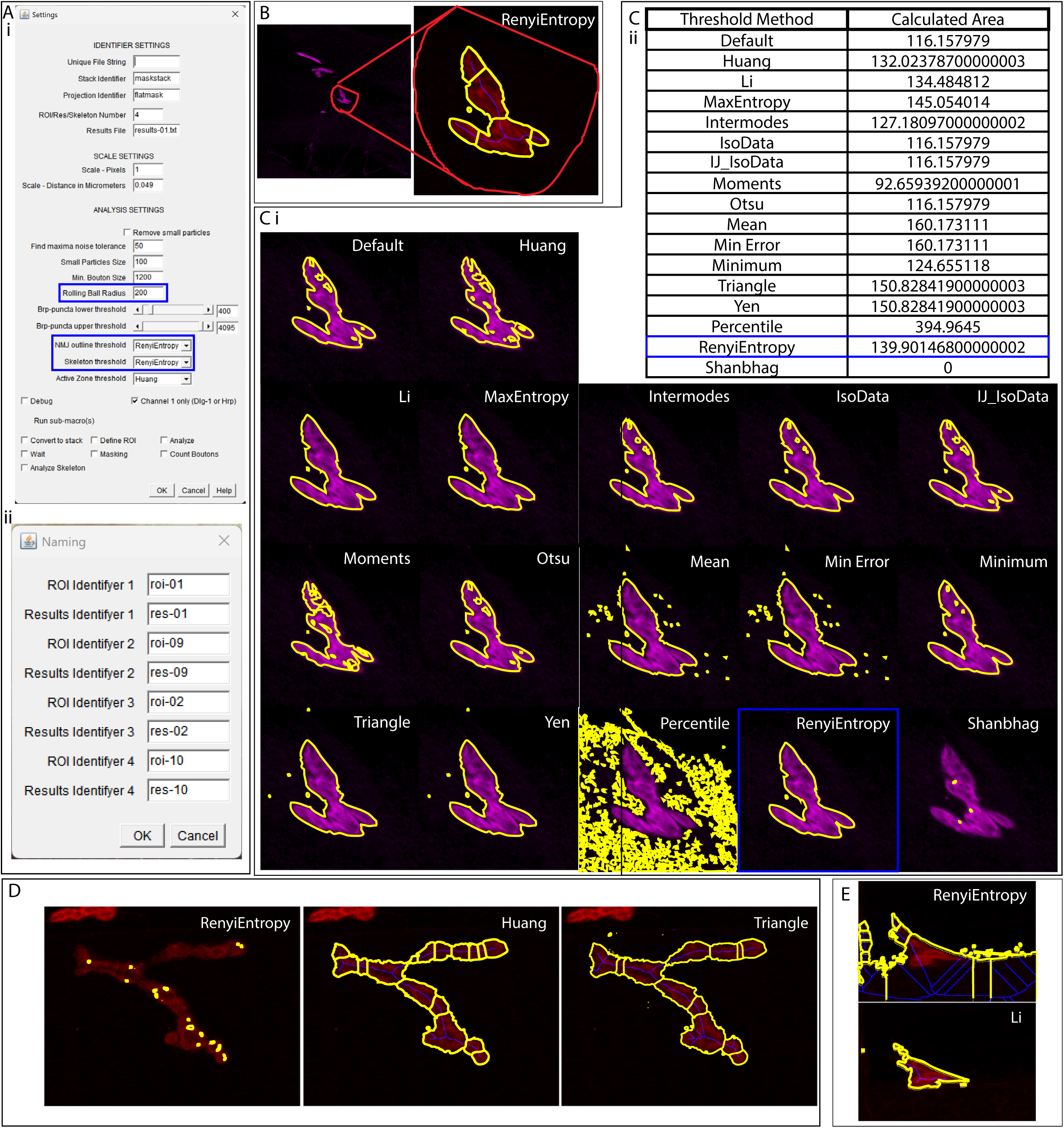
Threshold-based measurement of Dlg-positive NMJ area. (A) Screenshots of (i) the main Settings window, with the rolling-ball radius and threshold inputs highlighted, and (ii) the Naming dialogue box opened after selecting Analyze, used to assign the ROI and result identifiers. (B) Example of an appropriate NMJ outline generated with Rényi entropy thresholding. (C) Representative outlines (i) and calculated areas (ii) produced by Fiji thresholding methods. (D) When Rényi entropy excludes genuine Dlg-positive area, Huang or Triangle can produce a more complete outline. (E) When Rényi entropy includes background or non-NMJ signal, Li can produce a more restrictive outline.

**Note:** The parent quantification folder contains the text results file. Each analyzed hemisegment subfolder contains ROI-specific result images, including the generated outline and skeleton. Use a unique results filename because the plugin can overwrite an existing file with the same name.

**CRITICAL:** The thresholding method is an analysis decision. Apply a prespecified decision rule, blind the analyst to genotype when possible, and retain a record of any image-specific deviation from the default method.

### Count boutons and classify axon and NMJ branch points

#### Timing: approximately 8–10 min per image

This step manually counts boutons and distinguishes branch points that occur along Dlg-negative axon segments from branch points within Dlg-positive NMJ territory (**Figure 3A-G**).

1. Configure bouton and branch-point counting. **(Figure 3A)**
2. Open the NMJ Morphometrics plugin.
3. Set **Stack identifier** to maskstack and **Projection identifier** to flatmask.
4. Enter a results-file name, such as boutons_and_branchpoints.txt.
5. Select **Count Boutons**, set the four ROI identifiers, and select **Count Branching**.
6. Set the postsynapse identifier to flatmask and the axon identifier to GFP.
7. Select the parent quantification folder.
8. 6. Count axon branch points
9. Use the axon/Dlg composite to identify branch points located outside Dlg-positive territory.
10. Classify a branch point directly connected to the main axon stalk as first order, a branch point arising from a first-order branch as second order, and a branch point arising from a second-order branch as third order **(Figure 3B-D)**.
11. Enter the number of first-, second-, and third-order axon branch points in the dialog box.

**Figure 3.**
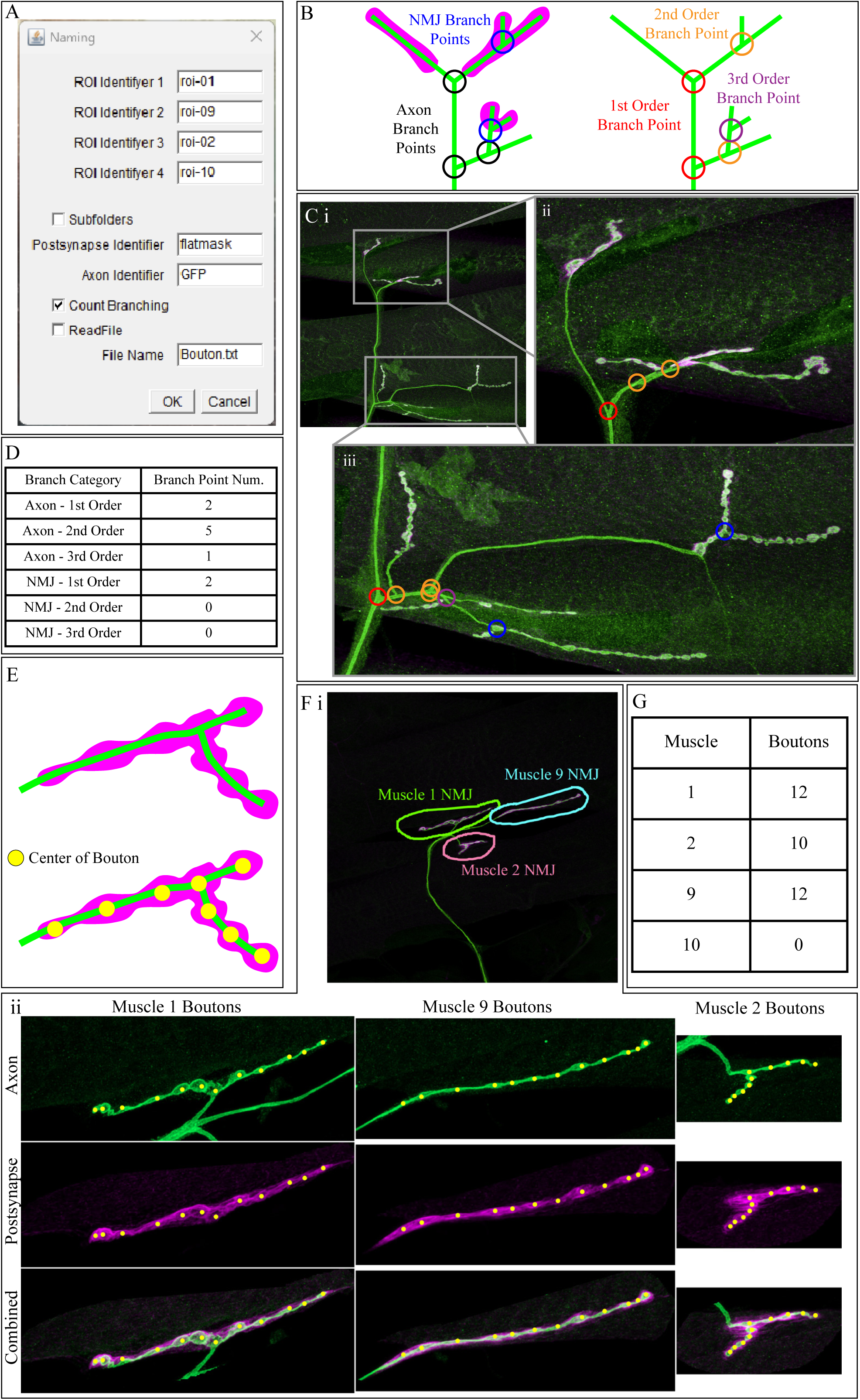
Counting branch points and boutons. (A) Screenshot of the Naming dialogue box opened after selecting Count Boutons, with Count Branching selected. (B) Branch points are classified as NMJ-associated when the bifurcation colocalizes with Dlg and as axonal when it lies outside Dlg-positive territory. Branch order indicates the number of branching levels from the main axon stalk. First-, second-, and third-order axonal branch points are indicated in red, orange, and purple, respectively; NMJ-associated branch points are indicated in blue. (C) Representative analysis of a phasic motor neuron showing the complete dorsal muscle field (i) and magnified regions (ii–iii). Colored circles identify classified branch points. (D) Branch-point counts from (C). (E) Schematic showing one point placed at the center of each bouton. (F) Muscle-specific ROIs (i) and bouton annotations (ii). (G) Bouton counts from (F).

**Note:** Classify a point according to the location of the bifurcation itself. Establish an exclusion rule before analysis for a branch point that cannot be assigned confidently because Dlg and axon signals overlap ambiguously.

7. Count boutons and NMJ branch points within each ROI
  a. Use the multipoint tool to place one point at the center of every morphologically distinct bouton in the axon/Dlg composite **(Figure 3E-G)**.
  b. Inspect all channels before finalizing the count. The macro records points placed in any channel, so mark each bouton only once.
  c. Count first-, second-, and third-order branch points whose bifurcations lie within Dlg-positive NMJ territory.
  d. Enter the branch-point counts and click **OK**.
  e. Repeat the analysis for every saved ROI.

**Note:** In the main results file, lines labeled with the axon filename contain first-, second-, and third-order axon branch counts; lines labeled with an ROI filename contain the bouton count and first-, second-, and third-order NMJ branch counts for that ROI. Each subfolder also contains the saved bouton-point file for each ROI.

**Note:** To review an existing analysis, select **Read File** and load the matching results file. Saved bouton points can then be inspected and corrected (Troubleshooting, problem 4).

8. Calculate the mean Dlg-positive area per bouton for each hemisegment
  a. Sum Dlg-positive area across all innervated muscle ROIs.
  b. Sum bouton counts across the same ROIs.
  c. Divide total Dlg-positive area by total bouton number.

**CRITICAL:** Report this value as mean Dlg-positive area per bouton. It is a two-dimensional proxy and is not a direct measurement of bouton volume.

### Clean the NMJ skeleton and quantify length

#### Timing: approximately 5 min per image

This step removes threshold-generated artifacts from the skeleton and reports the length of the Dlg-positive NMJ arbor (**Figure 4A-C**).

1. Generate and clean skeletons. (**Figure 4A**)
2. Open the NMJ Morphometrics plugin and set **Stack identifier** to maskstack and **Projection identifier** to flatmask.
3. Enter a results-file name, such as NMJ_skeleton_results.txt, and select **Analyze Skeleton**.
4. Enter res-01, res-02, res-09, and res-10 as the result identifiers and skeleton-01, skeleton-02, skeleton-09, and skeleton-10 as the corresponding skeleton identifiers.
5. Select the parent quantification folder.
6. Work only in the skeleton channel. Use the freehand-selection tool to delete spurious branches and small disconnected objects (**Figure 4B**).
7. Use a 3-pixel-wide pencil tool to reconnect a skeleton that fails to follow a genuine NMJ segment (**Figure 4C**).
8. Remove each artifact completely; a residual stub contributes to the calculated length.
9. Click **OK** to save the corrected skeleton and continue.

**Figure 4.**
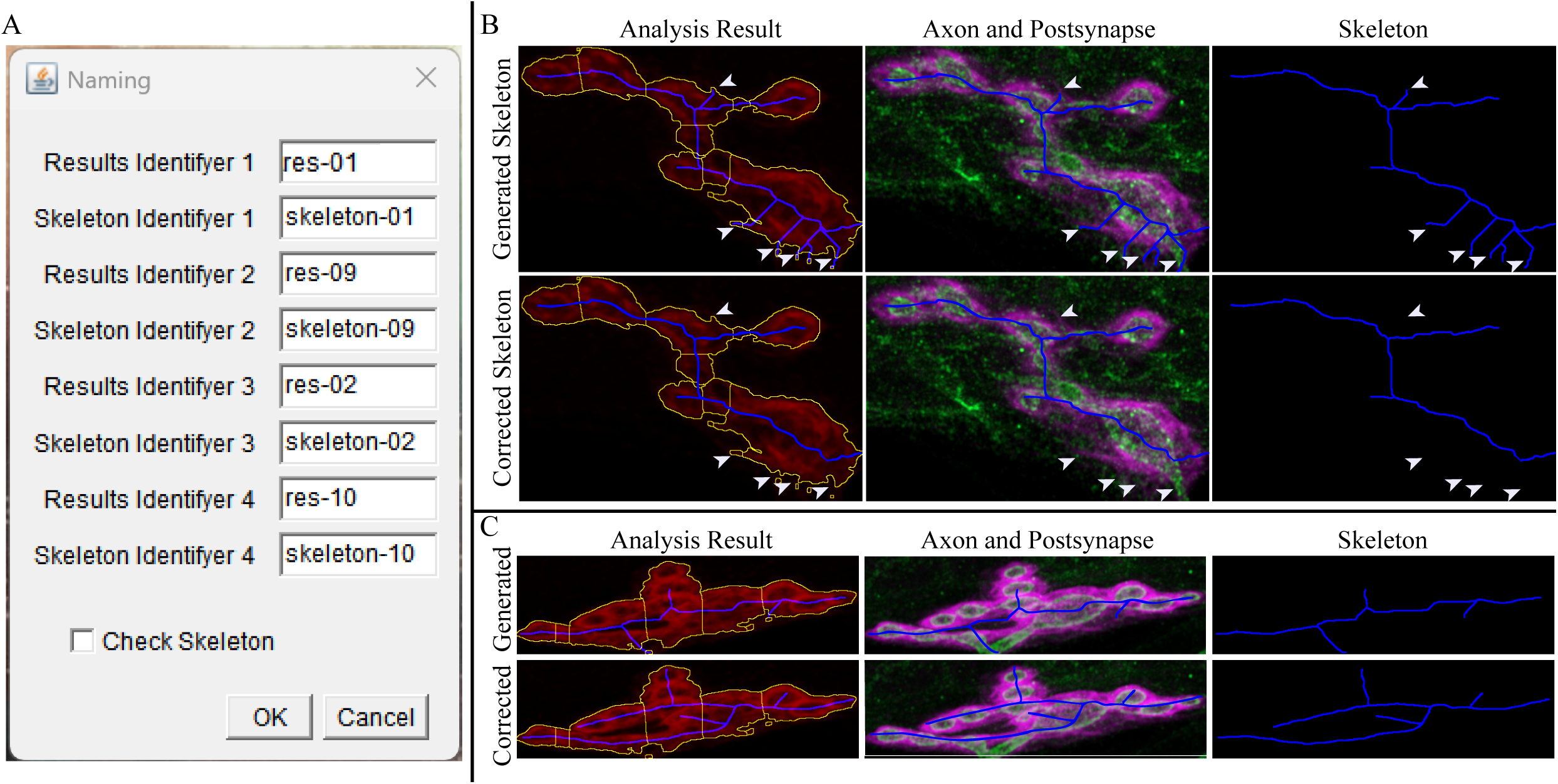
Correcting generated NMJ skeletons. (A) Screenshot of the Naming dialogue box opened after selecting Analyze Skeleton, showing the result and skeleton identifiers and the Check Skeleton option. (B) Threshold-generated skeletons can contain spurious branches. Arrowheads identify segments removed from the corrected skeleton. (C) When the skeleton does not follow a genuine NMJ segment, redraw the segment using the axon and Dlg channels as guides.

**Note:** Select **Check Skeleton** to reopen and verify previously corrected skeletons. Use the **Error** option to reload the current image without saving when an editing mistake occurs.

**Note:** The output includes total skeleton length, longest continuous path, branch number, and branch-point number for each ROI. Use total skeleton length as NMJ arbor length unless another output is prespecified.

### Quantify the whole-animal innervation pattern

#### Timing: approximately 20 min per larva

This step scores the native and ectopic targets of the labeled neuron across abdominal hemisegments and calculates muscle-specific innervation frequencies and phenotype penetrance (**Figure 5** and **Table 1**).

10. Acquire an overview image of the dorsal muscle field
  a. Use a 5x objective and enable Tiles and Z-stack in ZEN.
  b. Use Smart Setup to add the labeled-axon (e.g., Alexa Fluor 488), Dlg (e.g., Alexa Fluor 555), and muscle/phalloidin (e.g., Alexa Fluor 647) channels and establish nonsaturating settings.
  c. Select the confocal frame-size preset and use Tile Regions to define a field that contains abdominal segments A1–A6.
  d. Set the first and last z-positions to include the complete dorsal muscle field.
  e. Acquire the tiled z-stack and verify that the labeled axon, Dlg, and muscle boundaries can be resolved throughout the animal.

**Figure 5.**
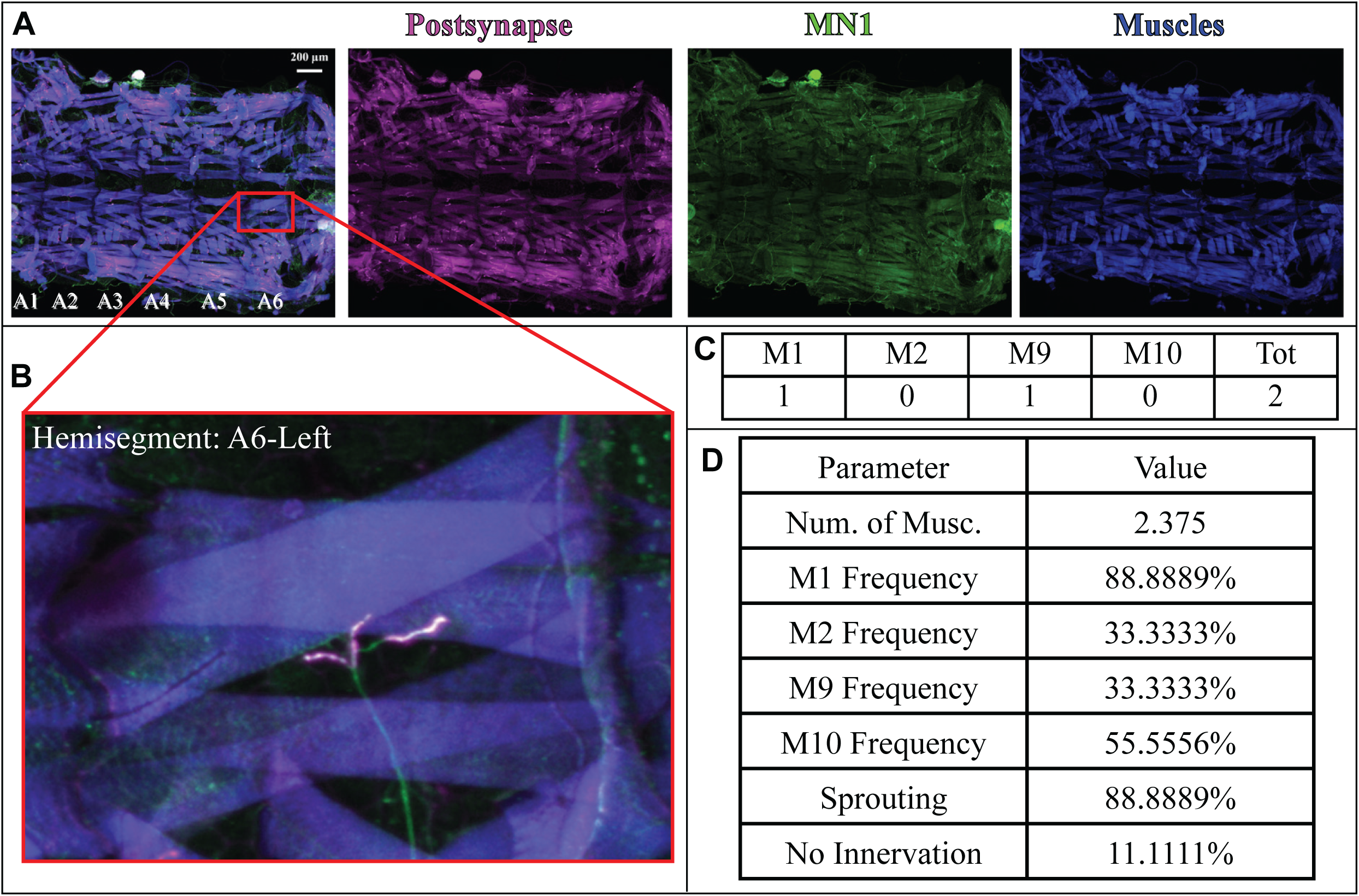
Quantifying the whole-animal innervation pattern. (A) Overview of abdominal segments A1–A6 in a larva after ablation of neighboring motor neurons. Muscles are blue, labeled MN1 is green, and Dlg is magenta. (B) Magnified A3-left hemisegment used to identify innervated muscles. (C) Binary scoring shows innervation of muscles 1 and 9 in the example hemisegment. (D) Per-animal summary of the mean number of muscles innervated, muscle-specific innervation frequencies, sprouting penetrance, and no-innervation penetrance.

**Table 1.** Example hemisegment-level scoring of the whole-animal innervation pattern. A value of 1 indicates colocalization of labeled MN1 and Dlg on the indicated muscle; 0 indicates absence of innervation; and n/a indicates a damaged or

| Hemisegment | Muscle 1 | Muscle 2 | Muscle 9 | Muscle 10 | No innervation | Total muscles |
| --- | --- | --- | --- | --- | --- | --- |
| A6-right | n/a | n/a | n/a | n/a | n/a | n/a |
| A6-left | 1 | 0 | 1 | 0 | no | 2 |
| A5-right | n/a | n/a | n/a | n/a | n/a | n/a |
| A5-left | 0 | 0 | 0 | 0 | yes | 0 |
| A4-right | n/a | n/a | n/a | n/a | n/a | n/a |
| A4-left | 1 | 1 | 0 | 1 | no | 3 |
| A3-right | n/a | n/a | n/a | n/a | n/a | n/a |
| A3-left | 1 | 0 | 0 | 1 | no | 2 |
| A2-right | 1 | 1 | 0 | 1 | no | 3 |
| A2-left | 1 | 0 | 1 | 1 | no | 3 |
| A1-right | 1 | 0 | 1 | 0 | no | 2 |
| A1-left | 1 | 1 | 0 | 1 | no | 3 |

**Alternative:** A light-sheet microscope can be used when it provides sufficient resolution to establish colocalization between the labeled axon and Dlg.^1^

11. Score innervation in each hemisegment
  a. Identify abdominal segments A1–A6 and the left and right hemisegments.
  b. For each hemisegment, score muscles 1, 2, 9, and 10 as innervated (1) only when the labeled axon and Dlg-positive postsynaptic structure colocalize on that muscle (**Figure 5A-C**).
  c. Score a hemisegment as no innervation only when phalloidin confirms that the muscles are intact and no labeled axon or Dlg-positive structure is present on any of the four muscles.
  d. Mark a torn, folded, missing, or otherwise uninterpretable hemisegment as n/a and exclude it from all denominators.
  e. Repeat the scoring for every analyzable hemisegment in each larva.
  12. Calculate whole-animal outcomes (**Figure 5D**)
    a. For the mean number of muscles innervated, count positive muscles in each innervated hemisegment and average across innervated hemisegments. Exclude damaged hemisegments and hemisegments scored as **no innervation** from this mean.
    b. For the innervation frequency of muscle *m*, divide the number of intact hemisegments in which muscle *m* is innervated by the total number of intact, analyzable hemisegments, including hemisegments scored as **no innervation**, and multiply by 100.
    c. For sprouting penetrance, divide the number of intact hemisegments with ectopic innervation of muscle 2, 9, or 10 by the total number of intact, analyzable hemisegments and multiply by 100.
    d. For no-innervation penetrance, divide the number of intact hemisegments scored as **no innervation** by the total number of intact, analyzable hemisegments and multiply by 100.

**CRITICAL:** Use the same denominator definition for every animal and condition. The muscle-frequency and phenotype-penetrance calculations include intact no-innervation hemisegments; only the mean number of muscles innervated excludes them.

## Expected outcomes

This workflow produces one masked Dlg stack and projection per hemisegment, muscle-specific ROIs, thresholded NMJ outlines, bouton annotations, cleaned skeletons, and tabulated outputs for Dlg-positive area, bouton number, mean Dlg-positive area per bouton, NMJ arbor length, and first-, second-, and third-order axon and NMJ branch points (Figure 6A-G). The whole-animal analysis produces a hemisegment-level innervation matrix and per-animal measurements of muscle-specific innervation frequency, sprouting penetrance, and no-innervation penetrance (Figure 6I).

**Figure 6.**
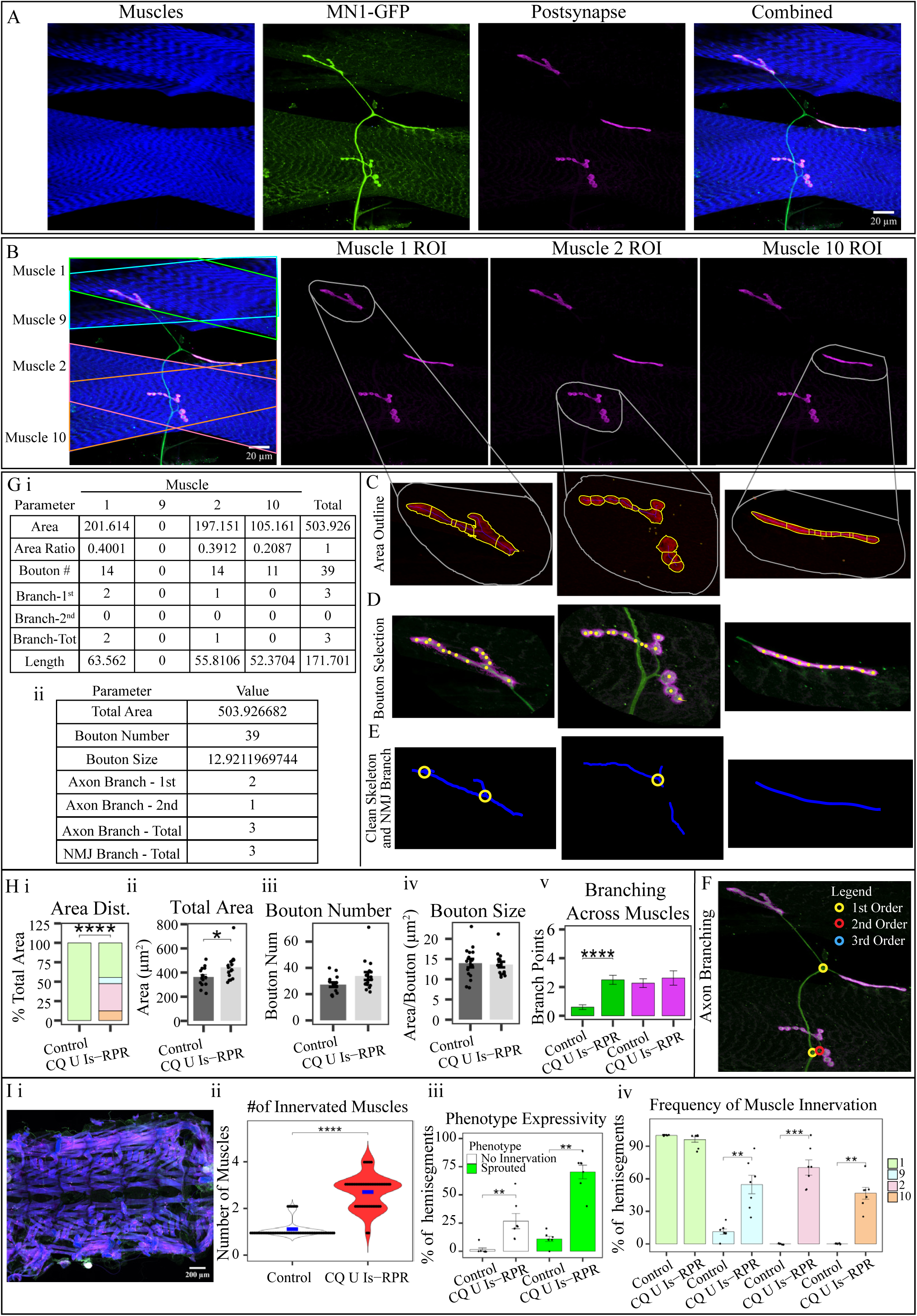
Representative outputs and expected results of compensatory sprouting quantification. (A) Representative hemisegment showing muscles (blue), labeled MN1 (green), Dlg-positive postsynaptic structures (magenta), and the merged image. (B) Muscle identities and muscle-specific ROIs for the Dlg-positive structures retained after masking; muscle 9 is not innervated in this example. (C) Threshold-derived Dlg-positive outlines for the innervated muscle ROIs identified in (B). (D) Bouton annotations, with yellow points placed near the center of each bouton. (E) Cleaned NMJ skeletons (blue) and first-order NMJ branch points (yellow circles). (F) Axonal branch points classified as first order (yellow), second order (red), or third order (blue) on the merged axon and Dlg image. (G) Example ROI-specific (i) and neuron-level (ii) outputs, including Dlg-positive area, muscle-specific area allocation, bouton number, mean Dlg-positive area per bouton, branch-point number, and NMJ arbor length. (H) Group summaries of muscle-specific area distribution (i), total Dlg-positive area (ii), bouton number (iii), mean Dlg-positive area per bouton (iv), and axon versus NMJ branch-point number (v). (I) Whole-animal innervation analysis showing a representative overview (i), mean number of muscles innervated (ii), sprouting and no-innervation penetrance (iii), and muscle-specific innervation frequency (iv). Data in panels H and I were published previously in Olsen et al.^1^

In the complete-denervation paradigm described by Olsen et al.^1^, MN1 retains its native NMJ on muscle 1 and frequently forms ectopic Dlg-positive NMJs on muscles 2, 9, and 10 (Figure 6A and 6B). Morphometric analysis can reveal an increase in total MN1 synaptic area and bouton number together with redistribution of synaptic area from the native target to ectopic targets (Figure 6G and Figure 6H, panels i-iv). An increase in Dlg-negative axon branch points without a corresponding increase in Dlg-positive NMJ branch points supports de novo axonal sprouting rather than simple expansion of established NMJ arbors (Figure 6E, Figure 6F, and Figure 6H, panel v). The exact magnitude and distribution of these outcomes depend on genotype, developmental timing, imaging quality, and sampling.

## Quantification and statistical analysis

Organize morphology data by hemisegment and retain a unique larva identifier so that measurements from the same larva can be identified. Calculate the following outcomes (Figure 6G):

• Total NMJ area = summed Dlg-positive area on muscles 1, 2, 9, and 10.

• Total bouton number = summed bouton counts on muscles 1, 2, 9, and 10.

• Mean Dlg-positive area per bouton = total NMJ area / total bouton number.

• Total axon or NMJ branch points = summed first-, second-, and third-order branch points in the corresponding category.

• Muscle-specific allocation = Dlg-positive area on muscle *m* / total NMJ area.

For an intact muscle with no MN1-associated NMJ, record muscle-specific area and bouton number as `0`. If the hemisegment has a nonzero total NMJ area, the allocation to an uninnervated muscle is also `0`. However, if the total NMJ area for the entire hemisegment is zero, muscle-specific allocation is undefined and should be recorded as `n/a`. Similarly, if total bouton number is zero, mean Dlg-positive area per bouton is undefined and should be recorded as `n/a`. Reserve `n/a` for undefined calculations and damaged or uninterpretable measurements.

Use the statistical tests described in the associated study^1^, treating the larva as the biological replicate and accounting for multiple hemisegments from the same larva. Report sample sizes, exact *p* values, and any multiple-comparison correction. Define image-quality and exclusion criteria before analysis.

## Limitations

This protocol requires strong, specific labeling of a single neuron. Multiple labeled axons can prevent reliable attribution of branches and Dlg-positive structures. Exclude hemisegments with off-target expression using prespecified criteria.

Measurements obtained from maximum-intensity projections are two-dimensional proxies and may merge structures that overlap in z. Dlg-positive area and mean area per bouton do not directly measure bouton volume, active-zone number, synaptic transmission, or muscle activity. Functional conclusions therefore require complementary assays.

Masking, ROI placement, thresholding, bouton and branch-point classification, and skeleton cleaning involve manual judgment and are sensitive to staining and imaging quality. Blind analysts when possible, apply consistent acquisition and processing settings, and archive intermediate files. Adapting the workflow to other neurons, developmental stages, or injury models requires appropriate validation.

## Troubleshooting

### Problem 1: The z-position shifts and part of the axon lies outside the acquired stack (before you begin, step 11)

#### Potential solution

Remove residual PBST from the forceps before transferring the fillet into Fluoromount-G; excess aqueous buffer dilutes the mounting medium and allows the specimen to move. Define a small z-margin above and below the visible axon. Exclude an image if any labeled axon or associated Dlg-positive structure is truncated rather than attempting to infer the missing morphology.

### Problem 2: The mask retains non-colocalized Dlg or removes genuine Dlg-positive signal (step-by-step method details, step 2)

#### Potential solution

Adjust the axon threshold to include the complete axonal profile and a narrow surrounding margin. If the restored mask selection includes non-colocalized Dlg, hold **Alt/Option** while using the freehand-selection tool to draw around and subtract the unwanted region. If associated Dlg is excluded, hold **Shift** while drawing around the missing region to add it to the selection (**Figure 1D-E**). Inspect the corrected selection before accepting the mask, and apply the same written decision rule across experimental groups. Do not use **Edit > Clear**, because this command changes the image pixels rather than only modifying the selection.

### Problem 3: The NMJ outline does not closely follow the Dlg-positive structure (step-by-step method details, step 4)

#### Potential solution

First verify that background subtraction is appropriate by previewing **Process > Subtract Background** on flatmask. Increase the rolling-ball radius if genuine boutons are removed. If Rényi entropy omits genuine NMJ area, test Huang or Triangle (**Figure 2D)**. If it includes background or non-NMJ signal, test Li (**Figure 2C and 2E**). Reanalyze only the affected ROIs, document the alternative method, and avoid choosing a method based on genotype or desired outcome.

### Problem 4: An incorrect bouton point cannot be removed after reopening the saved point file (step-by-step method details, step 7)

#### Potential solution

Activate the saved multipoint selection. With the freehand-selection tool, hold Alt/Option and draw around the incorrect point to subtract it from the selection. Confirm the corrected point count before resaving the point file. Do not use Edit > Clear, because that command changes image pixels.

### Problem 5: The masking or analysis routine does not start because the number of detected files differs among channels (before you begin, step 15; step-by-step method details, steps 1 and 4)

#### Potential solution

Confirm that every hemisegment folder contains one stack, one flatstack, and one matching GFP_ projection before masking, and one maskstack, one flatmask, and the expected ROI files before analysis. Remove duplicate exports, correct mismatched file-name prefixes, and ensure that identifiers entered in the dialog match the file names exactly. Avoid spaces and special characters in analysis identifiers.

### Problem 6: The generated skeleton contains spurious branches or fails to follow the NMJ (step-by-step method details, step 9)

#### Potential solution

Compare the skeleton with both the axon and Dlg channels. Delete disconnected objects and threshold-generated stubs completely. Redraw only segments that are clearly supported by the underlying signal using a 3-pixel-wide pencil tool (**Figure 4**). If extensive correction is required, return to the mask and thresholding steps because manual redrawing should not substitute for poor segmentation.

## Resource availability

### Lead contact

Further information and requests for resources and reagents should be directed to and will be fulfilled by the lead contact, Aref Zarin.

### Technical contact

Technical questions on executing this protocol should be directed to and will be answered by the technical contact, Lizzy Olsen.

### Materials availability

Fly lines generated and/or used in this study are available from the lead contact upon reasonable request.

### Data and code availability

Raw data and the MATLAB and R code used for analysis and visualization in the associated study are available through [Mendeley Data](https://data.mendeley.com/datasets/c9fdgs69xx/3). The original NMJ Morphometrics plugin is available at [Figshare](https://figshare.com/articles/dataset/Drosophila_NMJ_Morphometrics/2077399/1).

## Acknowledgments

We thank the Bloomington Drosophila Stock Center, Chris Doe, Troy Littleton, Dion Dickman, Keiko Hirono, and Sarah Ackerman for providing transgenic fly lines. We also thank Lauren Carlisle and Elaina Hildner for technical support. This study was supported by NIH-NINDS 1R01NS142268-01A1, the Texas A&M University Division of Research Targeted Proposal Teams (TPT) funding program, Zarin Ascend FY25 and FY26, and two internal grants from the Texas A&M College of Arts and Sciences, Zarin STRP FY24 and FY25.

## Author contributions

Conceptualization, L.O. and A.Z.; methodology, L.O. and A.Z.; software, L.O. and A.Z.; validation, L.O. and A.Z.; investigation, L.O. and A.Z.; formal analysis, L.O. and A.Z.; visualization, L.O. and A.Z.; writing - original draft, L.O. and A.Z.; writing - review and editing, L.O. and A.Z.; supervision, A.Z.; funding acquisition, A.Z.

